# Evolution of photochemical reaction centres: more twists?

**DOI:** 10.1101/502450

**Authors:** Tanai Cardona, A. William Rutherford

## Abstract

The earliest event recorded in the molecular evolution of photosynthesis is the structural and functional specialisation of Type I (ferredoxin-reducing) and Type II (quinone-reducing) reaction centres. Here we point out that the homodimeric Type I reaction centre of Heliobacteria has a Ca^2+^-binding site with a number of striking parallels to the Mn_4_CaO_5_ cluster of cyanobacterial Photosystem II. This structural parallels indicate that water oxidation chemistry originated at the divergence of Type I and Type II reaction centres. We suggests that this divergence was triggered by a structural rearrangement of a core transmembrane helix resulting in a shift of the redox potential of the electron donor side and electron acceptor side at the same time and in the same redox direction.

## Evolution of Photosystem II

There is no consensus on when and how oxygenic photosynthesis originated. Both the timing and the evolutionary mechanism are disputed. The timing ranges from the early Archean, 3.7 billion years ago [1–4] to immediately before the **Great Oxidation Event** (see Glossary), 2.4 billion years ago [5, 6]. Mechanisms proposed range from ancient gene duplication events involving the photosystems in **protocyanobacteria** [7, 8], to more recent horizontal gene transfer events into a non-photosynthetic ancestor of Cyanobacteria [9, 10]. However a complete scenario for the evolution of oxygenic photosynthesis should first explain how and when water oxidation to oxygen originated at the level of the **photochemical reaction centre**. That is, how and when Photosystem II (PSII) evolved the **Mn_4_CaO_5_ cluster** and the oxidising photochemistry required to split water (Figure 1).

In PSII the Mn_4_CaO_5_ cluster is coordinated by D1 and CP43 (Figure 2) [11]. The carboxylic C-terminus of A344 of D1 acts as a bidentate ligand that bridges the Ca^2+^ and a Mn atom [13], numbered Mn2 in Figure 2. Several Mn ligands, D342, E333, and H332 are also provided from the C-terminal domain of D1. In the same region, H337 provides a hydrogen-bond to a bridging oxygen (O3). In addition, two more ligands are provided from the inner part of D1, D170 and E189, located in the region connecting the 3^rd^ and 4^th^ transmembrane helices of D1. Furthermore, E354 from CP43 provides a bidentate ligand bridging to Mn2 and Mn3, and R357 provides a hydrogen-bond to O4 and is within 4.2 Å of the Ca^2+^. These two residues are located in an extrinsic protein domain between the 5^th^ and 6^th^ helices of CP43 that reaches into the electron donor-side of D1.

After **charge separation** the oxidised chlorophyll (P_D1^+^_) extracts an electron from the kinetically competent redox active tyrosine, D1-Y161, known as Y_Z_, forming the neutral tyrosyl radical, which in turn oxidises the Mn_4_CaO_5_ cluster. Y_Z_ is hydrogen-bonded to H190 and the electron transfer step to P_D1^+^_ is coupled to a movement of the hydroxyl proton to H190. There is no Mn_4_CaO_5_ cluster in D2 (Figure 2), however the tyrosine-histidine pair is conserved in a symmetrical position in this subunit too (D2-Y160 and D2-H189) giving rise to another redox active tyrosine, Y_D_ [14, 15]. The presence of strictly conserved redox active tyrosine residues in both D1 and D2 led to the suggestion that water oxidation originated in a homodimeric ancestral photosystem with primordial Mn clusters on each side of the reaction centre [16]. It was also predicted that structural evidence for the vestiges of a metal binding site on D2 would exist [16]. This prediction was confirmed once refined structures of PSII became available [11, 13] (Figure 2) and its evolutionary significance has been discussed in more detail recently [17, 18].

**Figure 1.**
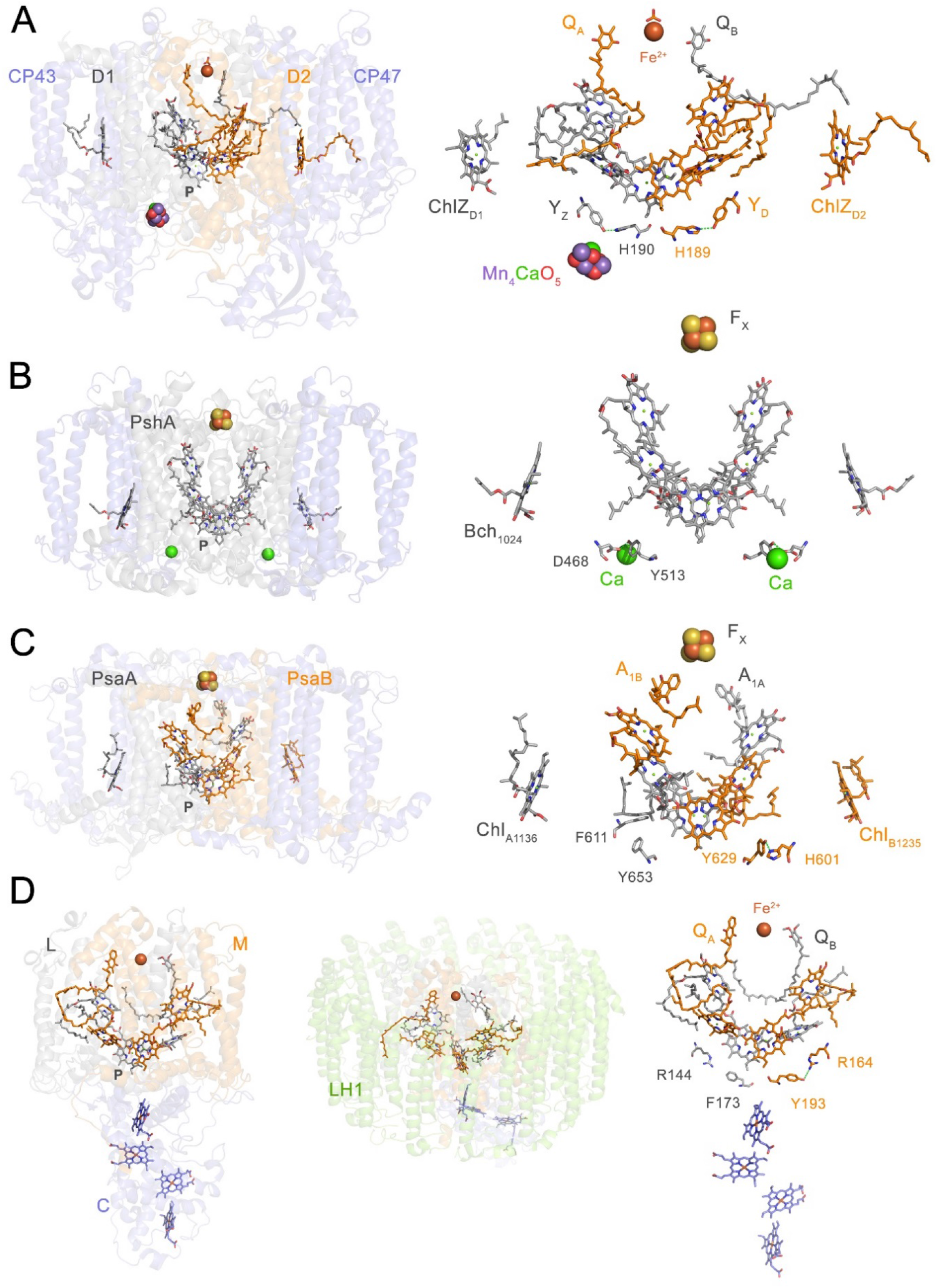
Photochemical reaction centres. Type II reaction centres function in quinone reduction and the last electron acceptor cofactor bound by the core reaction centre proteins is known as QB, an exhchangable quinone. Type I reaction centres function in ferredoxin reduction and the last electron acceptor cofactor bound by the core reaction centre proteins is known as FX, a Fe4S4 cluster. (**A**) PSII from the cyanobacterium *Thermosynechococcus vulcanus* (pdb id: 3wu2) [13]. Only the four main core subunits are shown: D1, D2, CP43 and CP47. The antenna subunits are highlighted in blue. On the right, a detailed view of the cofactors bound by the reaction centre proteins is shown. The Mn4CaO5 cluster is displayed as purple, red and green spheres. (**B**) Homodimeric Type I reaction centre from the phototrophic firmicute *Heliobacterium modesticaldum* (pdb id: 5v8k) [19]. Two Ca^2+^-binding sites with structural similarities to the Mn4CaO5 cluster are highlighted in green spheres. The reaction centre is made of a single protein, PshA, with the antenna domain highlighted in blue. (**C**) Photosystem I, the heterodimeric Type I reaction centre from the cyanobacterium *Thermosynechococcus elongatus* (pdb id: 1jb0) [20]. The reaction centre is made of two proteins, PsaA and PsaB, with the antenna domain highlighted in blue. (**D**) Anoxygenic Type II reaction centre from the phototrophic proteobacterium *Thermochromatium tepidum* (pdb id: 3wmm) [21]. Instead of a Mn4CaO5 cluster this reaction centre uses a bound cytrochrome (PufC) as direct electron donor to the photochemical pigments (P). Anoxygenic Type II reaction centre loss their ancestral antenna (highlighted in blue in **A, B**, and **C**) and evolved a new light harvesting complex (LH1). PSII is characterised by the presence of redox active tyrosine-histidine pairs. A hydrogen-bonded pair of residues at this position is likely an ancestral trait and can be found in all known reaction centres.

In the vicinity of Y_D_ in D2 there is no metal cluster and instead of ligands, several “space filling” phenylalanine residues are found (Figure 2). Like CP43, the CP47 subunit also has an extrinsic domain that reaches into D2, but instead of ligands, phenylalanine residues are also present, one of them located just 3.4 Å from Y_D_. These structural observations indicate that water oxidation could have originated in a homodimeric system before the duplication of the protein ancestral to D1 and D2, and that D2 evolved a unique hydrophobic space-filling plug to prevent the access of Mn and bulk water to Y_D_, thereby eliminating high potential catalysis on the D2 side of PSII. The evolutionary advantages of confining water oxidation to one side (the D1 side) of PSII have been discussed before in terms of the improved economy of having the highly oxidative chemistry confined to one side of the reaction centre, making frequent repair only necessary for D1 [22]. Other advantages include: 1) decreased energy losses and diminished photodamage compared to the situation when charge recombination can occur in the presence of two charge accumulating clusters per reaction centre [23], and 2) improved efficiency of photo-oxidative assembly of the Mn_4_CaO_5_ cluster [24].

The core of Type II reaction centres (D1 and D2 in PSII and L and M in anoxygenic Type II reaction centres) evolved from an ancestral homodimeric Type II reaction centre, but unlike the anoxygenic Type II reaction centres, PSII retained the CP43 and CP47 subunits, which originated from an ancestral homodimeric Type I reaction centre (Figure 1) [16, 25]. The current role of the CP43 in the ligation of the Mn_4_CaO_5_ cluster and CP47 in space-filling close to Y_D_, suggest an early function in cluster binding for the ancestral protein in the homodimeric forerunner of PSII. When taken with the fact that an *in series* Type II-Type I arrangement is only found in nature in oxygenic photosynthesis, these considerations led to the proposal that the evolution of the two types of reaction centre was linked to the origin of oxygenic photosynthesis itself [26].

Here we provide evidence supporting the premise that water oxidation originated at, or soon after, the divergence of Type I and Type II reaction centres, at one of the earliest stages in the evolutionary history of photosynthesis.

## A Ca^2+^-binding site in the Type I reaction centre from *Heliobacterium modesticaldum*

The recent structure of the homodimeric Type I reaction centre [19] from a basal anoxygenic photoheterotrophic firmicute, *Heliobacterium modesticaldum*, revealed a previously unknown Ca^2+^-binding site with a number of intriguing parallels to the Mn_4_CaO_5_ cluster of PSII (Figure 3). Firstly, this Ca^2+^-binding site is positioned at the electron donor side of each monomer of the reaction centre, in a location corresponding to that of the redox tyrosine-histidine pair in D1 and D2, in the immediate vicinity of the Mn_4_CaO_5_ cluster (Figure 3A). Secondly, the Ca^2+^-binding site is connected to the C-terminus of the reaction centre protein, PshA, by L605 and V608. L605 coordinates the Ca^2+^ via the backbone carbonyl and V608 via an oxygen from the C-terminal carboxylic group. This carboxylic acid ligand was not modelled in the structure [19], but can be seen in the electron density map (Figure 3E). In a similar way, the C-terminus of D1 provides direct ligands to the Mn_4_CaO_5_ cluster, i.e. D342 and A344 (Figire 3G). The A344, which is the last amino acid of the processed D1, is a ligand not only to Mn, but also to the Ca^2+^ via the carboxylic C-terminus. Thirdly, the Ca^2+^-binding site is linked to the antenna domain of PshA, via N263, which is located within the 5^th^ and 6^th^ transmembrane helices. N263 connects to the Ca^2+^ via two water molecules. In PSII, as described above, the CP43 residues E354 and R357, which are located in a large extrinsic domain between the 5^th^ and 6^th^ helices, connect the antenna to the Mn_4_CaO_5_ cluster; CP47 also shows a similar interaction with D2, but in this case contributing groups that block access to Y_D_ [18, 22]. Fourthly, D468, located at a position that overlaps with that of Y_Z_, provides a hydrogen bond to a tyrosine (Y513) found just 4 Å from the Ca^2+^. This resembles the Y_Z_-H190 pair, although PshA-Y513 and D1-H190 do not occupy identical structural positions (Figure 3C). In PSII, Y_Z_ is 4.7 Å from the Ca^2+^. The Y513-D468 hydrogen-bonded pair in the homodimeric reaction centre is found in the PsaB subunit of cyanobacterial PSI as Y629-H601 (Figure 3D), both of which are phenylalanine residues in PsaA.

In *H. modesticaldum*, it is not known whether this is a true Ca^2+^-binding site or the result of non-specific binding. None of the buffers used for purification of the complex [27, 28] or crystallisation conditions explicitly included Ca^2+^ salts [19], which may indicate that it is a binding site with significant affinity for Ca^2+^. Sequence comparisons suggests that a similar site might exist in homodimeric Type I reaction centres from the Chlorobi (Box 1). The presence of a divalent metal site in homodimeric Type I reaction centres so distant from PSII, yet in a manner so similar to the Mn_4_CaO_5_ cluster, is intriguing and potentially significant from an evolutionary perspective.

Structural parallels between the Ca^2+^-binding site in the homodimeric Type I reaction centre and the Mn_4_CaO_5_ cluster in PSII indicate that their most recent common ancestor had a site that was readily accessible to divalent cations at the donor side and near the photochemical pigments (P). The most recent common ancestor of homodimeric Type I reaction centres and PSII is the ancestral photosystem existing before the divergence of Type I and Type II reaction centres. This would mean that several of the structural elements needed for the origin of a water-oxidising cluster were already in place at this point in time: one of the earliest stages in the evolution of photosynthesis. These elements include the location itself, at least two ligands available from the C-terminus, at least one ligand available from the antenna domain, Ca^2+^, and even potentially the tyrosine. These structural similarities provide a rationale for the evolutionary origin, location and ligand sphere of the Mn_4_CaO_5_ cluster. They indicate a more ancient origin for the involvement of Type I-like core antenna proteins in the binding of the cluster than has been considered until now [29, 30]. It seems quite likely that the CP43 and CP47 subunits were retained in PSII, at least in part, because of their interaction with the electron donor site and their role in binding the cluster in the ancestral homodimer. The loss of the core antenna of anoxygenic Type II reaction centres would thus be linked to structural changes at the electron donor side associated with the evolutionary bifurcation towards the branch leading to L and M (characterised by the photochemical oxidation of cytochromes) and away from the branch leading to D1 and D2 (characterised by photochemical water oxidation) [31].

**Figure 2.**
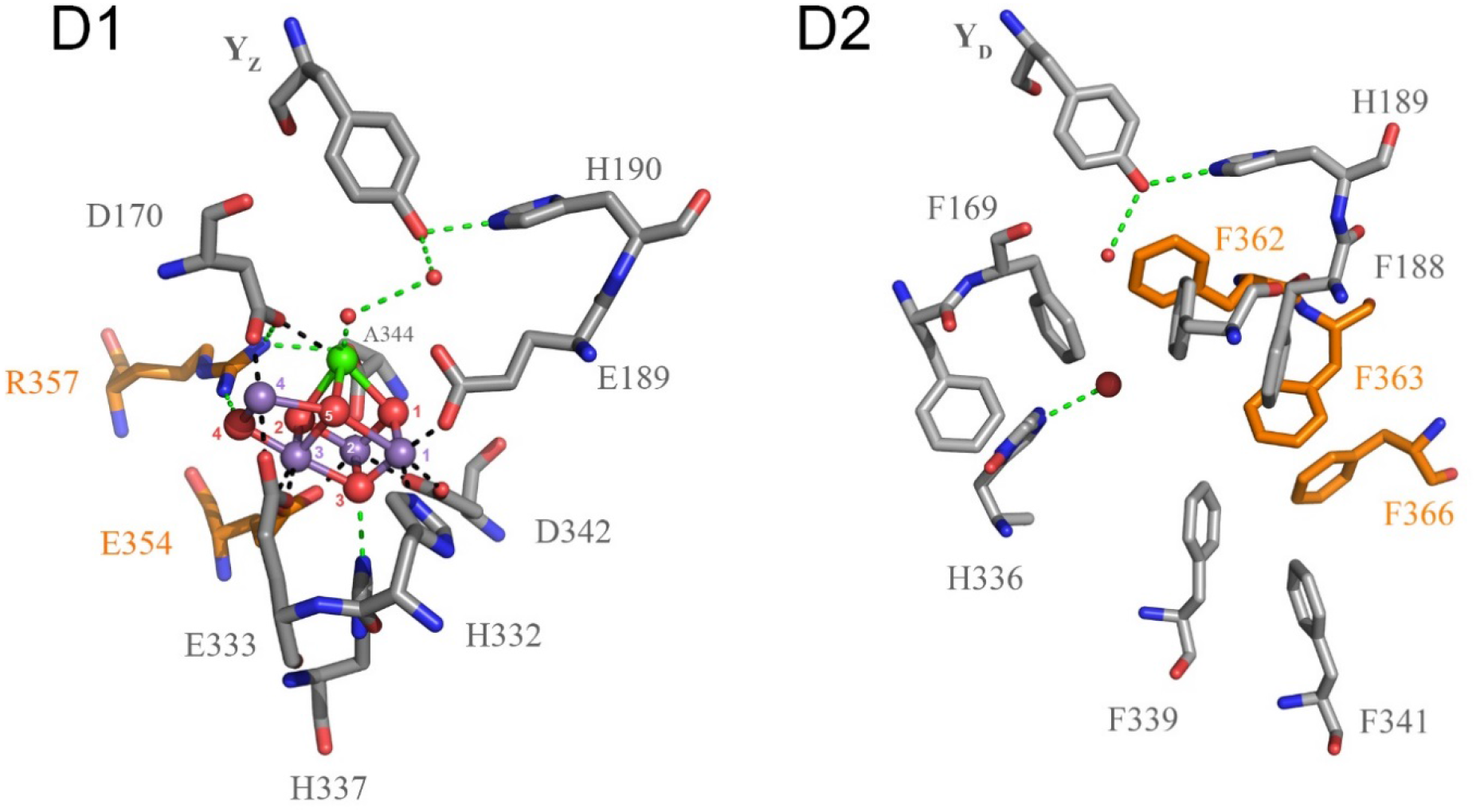
The Mn4CaO5 cluster of PSII. The cluster is coordinated by ligands from D1 (grey sticks) and CP43 (orange sticks). Ca^2+^ is connected to the redox tyrosine YZ (Y161) via hydrogen-bonded water molecules. On the right, the homologous site in D2 is shown. Residues from D2 are shown as grey sticks and those from CP47 as orange sticks. No catalytic cluster is observed but a strictly conserved redox tyrosine is found, YD (Y160). D2-F169 and D2-F188 occupy the positions of D1-D170 and D1-E189 respectively; and D2-F339 and D2-F341 occupy positions similar to those of D1-D342 and D1-A344, respectively. In addition, D2-H336 provides a hydrogen bond to a water molecule found between the phenylalanine residues in a way that is quite similar to the hydrogen bond of D1-H337 to the cubane oxygen (O3). CP47-F362 occupies the position of one of the waters that provides a hydrogen-bond to the phenolic oxygen of the YZ and which is thought to be important for its rapid, reversible, high-potential redox chemistry [32]. It has been suggested that water oxidation started in a homodimeric photosystem, with two catalytic clusters placed symmetrically, one on each side of the reaction centre and with ligands to the ancestral antenna protein [16, 18, 22].

**Figure 3.**
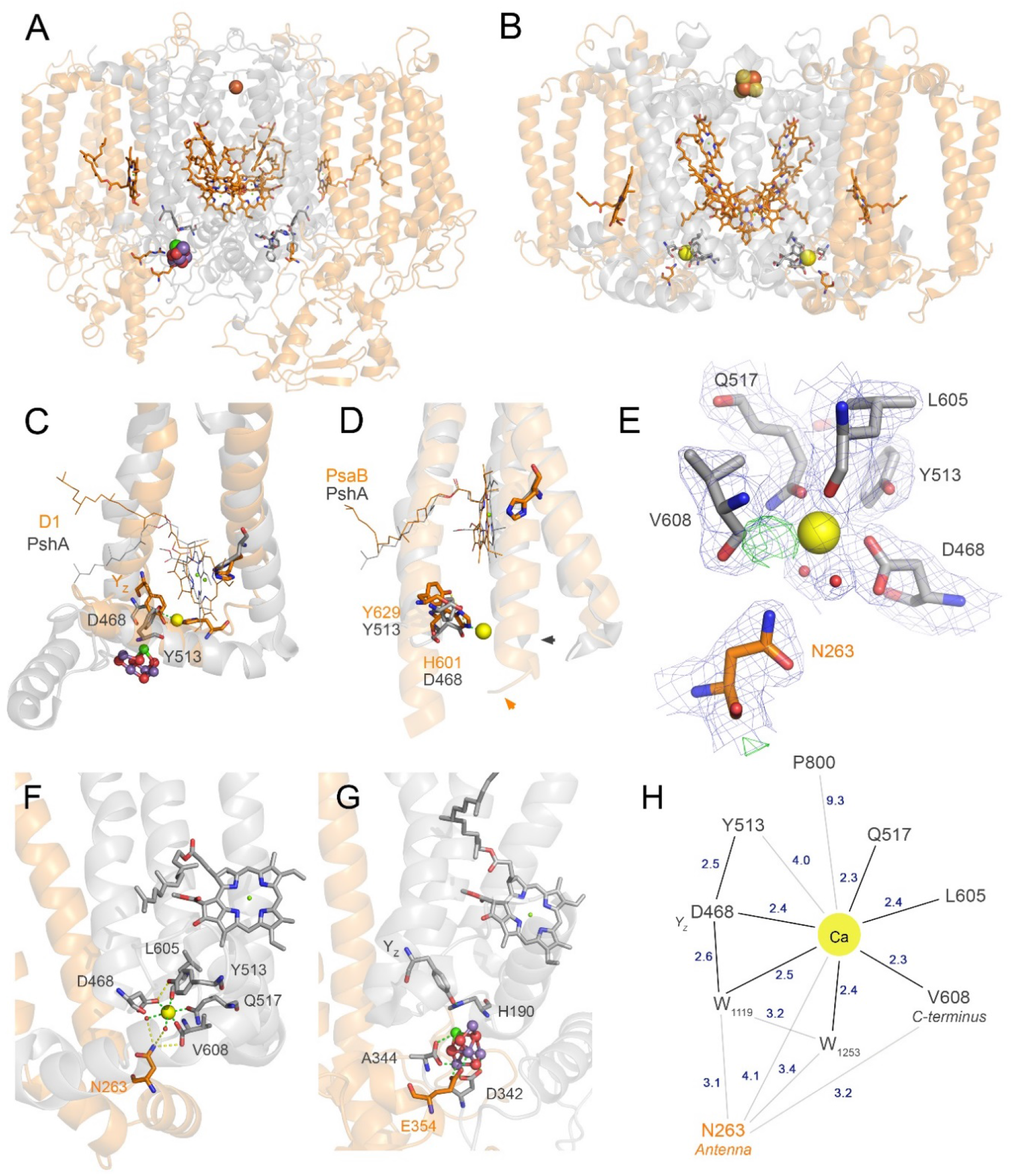
Structural parallels between the Mn4CaO5 cluster of PSII and the Ca^2+^-binding site of the homodimeric Type I reaction centre. (**A**) Full view of PSII showing in grey transparent ribbons the core proteins and in orange the antenna proteins. (**B**) Full view of the homodimeric Type I reaction centre from *Heliobacterium modesticaldum* showing in transparent grey the core domain and in orange the antenna domain of PshA. The yellow spheres are the Ca^2+^ ions located symmetrically on each side of the Type I reaction centre. (**C**) Overlap of D1 (orange) and PshA (grey). In PshA, Ca^2+^ is coordinated by D468, which occupies a structurally position similar to YZ in D1. Only the 9^th^ and 10^th^ transmembrane helices are displayed for clarity (3^rd^ and 4^th^ in D1). (**D**) Overlap of PsaB of cyanobacterial Photosystem I (orange), and PshA (grey). There is no Ca^2+^-binding site in Photosystem I as the 11^th^ transmembrane helix of the core domain is about two turns longer (arrows). The arrows mark the C-terminus. Only the last three transmembrane helices are shown and the interconnecting loops were hidden for clarity. (**E**) Electron density map of 5v8k around the Ca^2+^-binding site (blue mesh, contour map: 1σ). V608, the C-terminus carboxylic group, was modelled in the released structure as a carbonyl, missing the oxygen that coordinates Ca^2+^ [19]. The difference between the observed and calculated maps shows the positive density between V608 and the Ca^2+^ corresponding to the missing coordinating oxygen bond (green mesh, contour map: 3σ). (**F**) Close-up of the Ca^2+^-binding site showing the connection to the antenna domain via N263 and the C-terminus. (**G**) Close-up of the Mn4CaO5 cluster highlighting the connection to the antenna via E354. A344, the C-terminus, provides a direct ligand to Ca^2+^ in PSII. (**H**) Scheme of the Ca^2+^-binding site of PshA showing the closest distances (Å) to residues in the immediate vicinity. W stands for water and the words in italics highlight structural parallels to PSII.

### Box 1. Do other homodimeric Type I reaction centres have a Ca^2+^-binding site?

Sequence alignments of PscA and PshA indicate that the Type I reaction centre of phototrophic Chlorobi should retain a Ca^2+^-binding site like that found in *Heliobacterium modesticaldum*. The predicted Ca^2+^-binding site in the homodimeric Type I reaction centre of Chlorobi would be coordinated by the C-terminal A731 and by the backbone carbonyl of L729. Residue Y513 in the reaction centre of *H. modesticaldum* is conserved in all Chlorobi as Y599 (numbering from PscA of *Chlorobium limicola*). Residue D468 in *H. modesticaldum* is also conserved as D563 in Chlorobi. Q517 is not conserved in Chlorobi, instead a glutamate residue is found in this position, which can also act as a ligand. The loop region between the 5^th^ and 6^th^ transmembrane helices in the PscA antenna domain is larger than that found in PshA, but smaller than in CP43 and CP47. Due to big structural differences in this extrinsic loop, a residue equivalent to N263 cannot be identified in the Chlorobi sequence, instead we suggests that in Chlorobi this region will bind the Ca^2+^ via a strictly conserved glutamate at position 323 (or alternatively at position 345). Crosslinking experiments in the reaction centre of *Chlorobaculum tepidum* showed that K315 and K338, located in the extrinsic domain between the 5^th^ and 6^th^ transmembrane helices of the antenna domain, bound the heme-containing region of PscC, the immediate electron donor to P [12]. This is consistent with the antenna domain interacting with the electron donor site of the reaction centre as it is the case in the reaction centre of *H. modesticaldum* and PSII. It indicates that the Mn_4_CaO_5_ cluster evolved at the ancestral electron donor site like that found in homodimeric Type I reaction centres. Therefore, the position of PufC (the tetraheme cytochrome of anoxygenic Type II reaction centres, see Figure 1) and its evolution as an electron donor, represent a novel adaptation rather than the primitive ancestral state of Type II reaction centres.

## The divergence of Type I and Type II reaction centres

One of the earliest events in the evolution of photosynthesis was the structural and functional specialisation that produced the two known types of reaction centres. Structural comparisons suggest that major structural modifications had to occur at the divergence between the two types (Figure 4). Of particular importance are the changes that determined 1) the nature of the electron donor, 2) the nature of the electron acceptor, 3) the position of the ligand to P, and 4) the splitting (or fusion) of the core antenna and the reaction centre core.

The nature of the terminal electron acceptor of the ancestral reaction centre before the divergence of Type I and Type II is not known [31] (see outstanding questions). Various arguments have been given in favour of the most recent common ancestor of reaction centres reducing either iron-sulfur proteins [8, 33, 34] or alternatively, quinones [29, 35] as the terminal electron acceptor. Available phylogenetic and structural data however are inconclusive. Here, in line with earlier arguments [16, 36] and to illustrate the structural changes at the divergence of Type I and Type II reaction centres, we assume that the ancestor was more “Type I-like”, in that it reduced a soluble ferredoxin-type iron-sulfur protein and had a central core made up of a homodimer with each subunit containing 11 transmembrane helices. We also assume that it had an F_X_-like Fe_4_S_4_ cluster connecting the homodimeric core subunits and that the subunit containing the FAF_B_ clusters was soluble and exchangeable, as it appears to be in Heliobacteria [37, 38].

### Increasing the potential of P^+^

Previously much of the focus of discussion was on the changes that were responsible for making a reaction centre capable of oxidising water [22, 39, 40]. The E_*m*_ for water oxidation to O_2_ is +820 mV at pH 7 [41]. However the concentration of oxygen in the Archean atmosphere was likely well below 10^-5^ of the present level [42, 43], which translates to a concentration of dissolved oxygen in water below 2 nM [44]. Under these conditions, the E_*m*_ of water oxidation to oxygen is +720 mV [44]. The E_*m*_ of chlorophyll *a* in dichloromethane is +800 mV [40, 45], similar to that for water oxidation, but more driving force is needed to achieve efficient catalysis. In PSII the Mn_4_CaO_5_ cluster acts as the active site and the accumulator of oxidising equivalents in the form of higher valence Mn ions, in close association with Y_Z_, which serves as an electron relay and an electrostatic trigger. This catalytic system is oxidised by the photochemical pigments, known in PSII as P680. In present day PSII, the E_*m*_ of P680^+^/P680 is estimated to be about +1200 mV [46, 47]: 400 mV more oxidising than chlorophyll *a* in an organic solvent. The E_*m*_ of P700^+^/P700 is estimated to be about +450 mV [48, 49]: 350 mV less oxidising than chlorophyll *a* in an organic solvent. This makes the difference between the E_*m*_ of P680^+^/P680 and P700^+^/P700 about 750 mV and indicates that the protein environment strongly modulates the E_m_ of the photochemical pigments in both directions and in both types of reaction centre relative to that of the isolated pigment [40].

The idea that these differences in redox potential could have been achieved by only a limited number of mutations, an *oxidative jump*, rather than a very gradual shift to higher potential, was proposed when it was realised that this jump likely occurred in homodimeric reaction centres, since each mutation in the single gene could have a double effect in the homodimer [16, 22]. The nature of the likely changes were discussed [16, 22], but the factors responsible for the increased potential became clearer when computational approaches were applied with the advent of refined crystal structures [40, 50]. The main changes responsible were electrostatic, including significant effects from the charges of the metal cluster, the protein backbone dipoles from the ends of the transmembrane helices, and specific amino acid side chains. In fact, Ishikita *et al*. [40] calculated that in the absence of any protein charges or specific electrostatic effects the E_*m*_ of P in PSI and PSII would be closer to that of chlorophyll *a* in an organic solvent, about 720 mV for both systems.

The problem with the *oxidative jump* scenario in the evolution of PSII was that it failed to address a basic energy problem. Both PSII and PSI work on chlorophyll *a* photochemistry and, as such, are essentially the same colour and thus have the same energy available. Thus an upshift in the oxidising potential of the donor-side must be accompanied by a matching upshift in the potential of the acceptor-side. Without this, the reducing power of the excited state P* would not be capable of reducing the very low potential electron acceptors, typical of PSI, which are assumed to have been present in the homodimeric ancestor. The lack of a matching increase in the potential of the electron acceptors would have resulted in the loss of charge separation or at least a marked decreases in the quantum yield. This is exactly what happened when anoxygenic Type II reaction centres were engineered to increase the oxidising power of P^+^/P without increasing the potential of the electron acceptors [51]. Consequently, this oxidative jump scenario is faced with the problem that increasing the potential of the P^+^/P will only be feasible when the acceptor side is already oxidising. This then requires the homodimeric ancestor to have evolved a high potential acceptor side without any obvious evolutionary advantage or selection pressure. A solution to this problem would be that the acceptor side also underwent an oxidative jump at the same time as the donor side. Just such a dual effect might result from a helix-shift in the reaction centre core. A comparison of the crystal structures shows evidence for such a helix shift, see below and Figure 4.

### Increasing the potential of the acceptor side

The > 500 mV difference in the potential of the A_1_ phylloquinone (about –700mV) and the Q_A_ plastoquinone (−150 mV) is attributed to 1) a ∼500 mV electrostatic difference due to the net negative charge on F_X_ compared to the net positive charge on the non-heme iron [52], 2) the intrinsic redox potential difference between phylloquinone and plastoquinone (180mV for the n = 2 couples and ∼100 mV for the 1-electron reduction without protonation), and 3) the electrostatic environment provided by the protein [50, 53]. Therefore, a crucial event occurring at the divergence of Type I and Type II reaction centres is the swap of FX for the non-heme Fe^2+^. The ligands for F_X_ are two cysteines on a loop between the 7^th^ and 8^th^ transmembrane helices (2^nd^ and 3^rd^ in Type II reaction centres), see Figure 4A. These cysteines are not in positions homologous to those for the non-heme Fe^2+^, which arise from histidine residues near the top of the transmembrane helices equivalent to the 10^th^ and 11^th^ helices in Type I reaction centres (4^th^ and 5^th^ in Type II), see Figure 4C and D. F_X_ itself is in a position on the same central axis and closer to the membrane surface than the non-heme Fe^2+^. The quinones are located in similar but non-identical positions, with the A_1_ phylloquinones slightly closer to P and with their O-O axis tilted ∼60° relative to the membrane plane [20, 54]. Only the distal carbonyls of A_1_ are hydrogen-bonded to a peptide-bond nitrogen [20], which is in contrast to the situation for Q_A_ and Q_B_, where both carbonyls are hydrogen-bonded. In Type II reaction centres the quinones are thus held near parallel to the membrane [11, 55]. In the homodimeric Type I reaction centre structure no bound quinones were observed [19], but they have been detected by other methods [27, 56–59] and are thus likely to be loosely or transiently bound. A swap from F_X_ to a non-heme Fe^2+^/quinone system must have required drastic structural changes. Given the symmetry of the existing structures it seems likely that these must have occurred in a homodimeric ancestor, but even so it is not easy to predict the order of events or even the nature of the events that would have led to such a crucial change. Some insight however might be gained from the structural data available.

**Figure 4.**
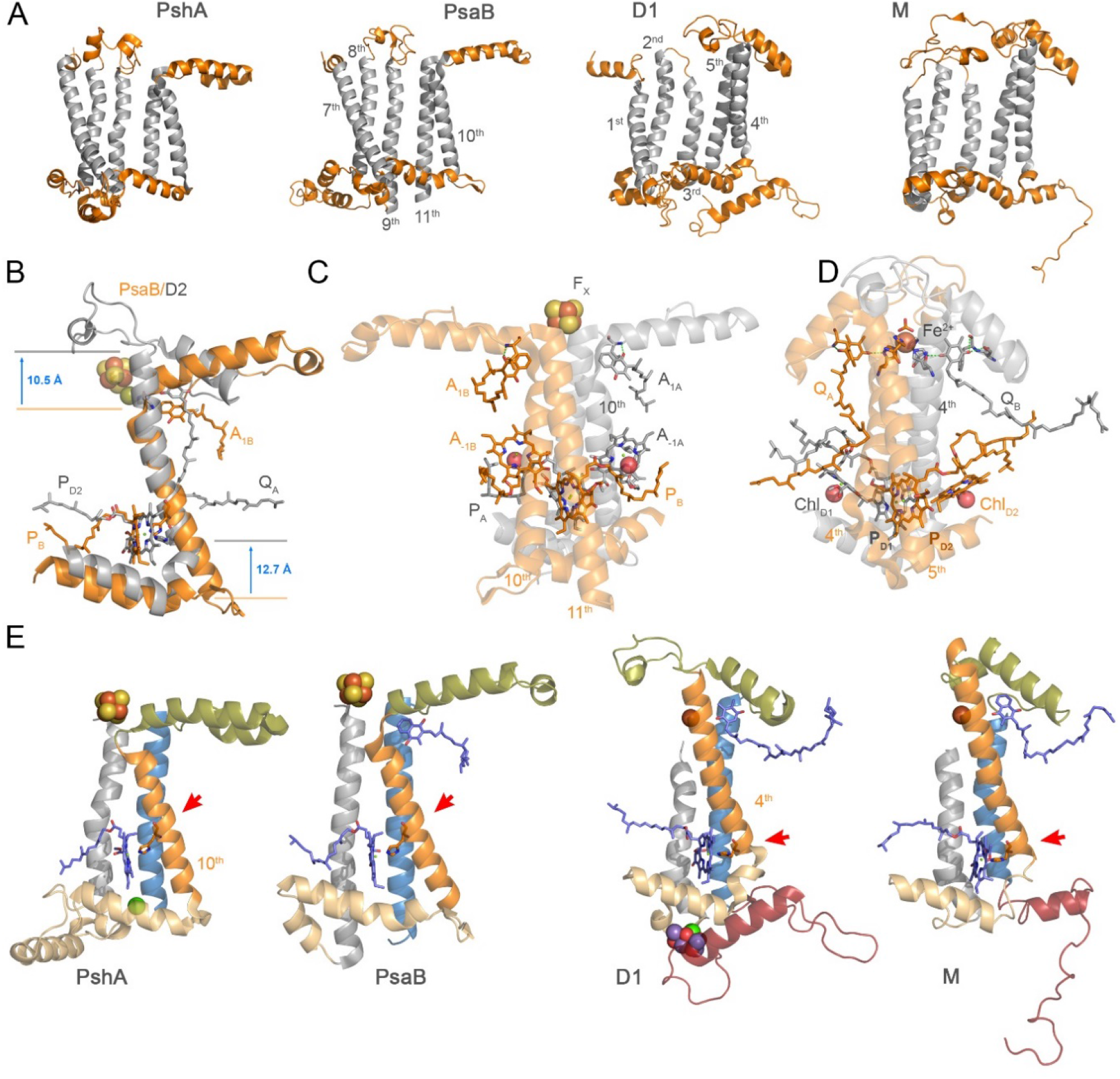
Structural comparisons of the reaction centre proteins. (**A**) PshA and PsaB exemplify Type I reaction centre proteins (core domain only) and D1 and M exemplify Type II reaction centre proteins. The transmembrane helices have been coloured grey and the interconnecting loops in orange. In Type I reaction centres, between the 8^th^ and 9^th^ helices there is a small loop that provides coordination to FX. This loop is absent in Type II (2^nd^ and 3^rd^ helices). (**B**) Overlap of PsaB (orange) and D2 (grey): only the helix that provides the ligand to P is shown (10^th^ in Type I, 4^th^ in Type II). The start and the end of the 4^th^ transmembrane helix in Type II reaction centres are shifted 12.7 Å and 10.5 Å respectively, relative to Type I. The position of the P chlorophylls remains unchanged. (**C**) A view of the phylloquinone binding sites in Photosystem I relative to the position of P and A-1. Only the 10^th^ and 11^th^ helices are shown. The red spheres represent the coordinating waters of A-1. The A0 electron acceptor chlorophylls were omitted for clarity. (**D**) A view of the plastoquinone binding sites in PSII relative to the position of P and ChlD1/D2. The red spheres represent the coordinating waters of ChlD1/D2. The pheophytin electron acceptors, the pigments at homologous positions to A0 in Type I, were omitted for clarity. (**E**) The last three transmembrane helices of the core reaction centre proteins are compared in order to highlight the structural differences between the two types. The red arrows mark the position of the histidine ligand to P. FX is 2.8 Å closer to P in the reaction centre of *H. modeticaldum* than in Photosystem I. All structures have been aligned to the position of the Mg atom of P and maintaining the last transmembrane helix vertical (blue helix).

### A change of position of the 10^th^ transmembrane helix

In Type I reaction centres the histidine ligand to P is in the middle of the transmembrane helix (Figure 4A and E), while in Type II reaction centres the histidine ligand is at the bottom of the helix. Despite this difference in the location of the liganding histidine residues, the P chlorophylls are at the same positions relative to the membrane plane, with the histidine in Type I reaching downwards, while the histidine in Type II is at the same level as the Mg^2+^.

This difference indicates that a change in the position of the helix took place. It required an upward movement of about 12 Å relative to a Type I reaction centre (Figure 4B and E), while the position of P remained unchanged. The change in position of the helix would have altered the interconnecting domains between the 9^th^ and 10^th^ helices and between the 10^th^ and 11^th^ helices, which make part of the donor and acceptor side, respectively. Such a shift of the position of the helix within the membrane would have changed the electrostatic environment of the photochemical pigments and likely affected any existing tuning of the redox potentials. Furthermore, the shift probably disrupted the assembly of F_X_ and would have inevitably changed groups on the electron acceptor end of the same helix. It seems possible that a histidine, near the top of the 10^th^ helix, could have become the ligand for the non-heme Fe^2+^. The homodimeric nature of this change meant two ligands were provided to the iron in one event. This transition should have favoured the selection of the second histidine from the 11^th^ helix (5^th^ helix in Type II reaction centres) to provide a strong stable central symmetrical Fe^2+^ coordination sphere, leaving two coordination positions in the Fe^2+^ empty to be filled by exchangeable ligands (water or by bicarbonate) as still exists in PSII today.

The newly formed His-Fe^2+^-His motif could have provided hydrogen-bonds to the distal carbonyl of available quinones, making their O-O axes near parallel with the membrane, and providing the quinone-quinone electron transfer pathway with the Q-His-Fe^2+^-His-Q motif that is characteristic of Type II reaction centres. The tight binding of the non-heme Fe^2+^ at the top of the transmembrane helices, holding the two monomers together, would have made the structural role of F_X_ redundant and could have hindered the folding of the loop that provided the cysteine ligands leading to its loss in Type II reaction centres.

The difference in the position of P relative to the transmembrane helix resulted in an up-shift in the E_*m*_ of P of Type II reaction centres of at least +140 mV, but possibly more. This +140 mV up-shift in potential relative to Type I is caused by the decrease in the effect of the protein backbone dipoles at the base of the transmembrane helix due to the different positions of the histidine ligand to P [40]. It may appear that the structural changes described above could not account for the large increase in the potential of P required for water oxidation. But, if the ancestral reaction centre was Type I-like, the electrostatic effects responsible for down-shifting the potential of P relative to the intrinsic chlorophyll *a* potential (+720–800 mV) would have been displaced after the structural change associated with the helix shift. This could account for an increase of up to +380 mV [40]. Altogether, the structural rearrangements could easily account for an increase in the E_m_ of P that could have kick-started water oxidation in anaerobic conditions upon a single structural change.

In addition, PSII has in common with Type I reaction centres not only the presence of antenna (CP43 and CP47) and the core peripheral chlorophylls (ChlZ_D1_/ChlZ_D2_), but also the fact that the monomeric chlorophylls (A_-1_ in Type I and Chl_D1_/Chl_D2_ in PSII) are coordinated by water molecules (Figure 4C and D). In anoxygneic Type II reaction centres, the Mg^2+^ of the equivalent bacteriochlorophylls are coordinated by histidine ligands, indicating that the absence of these histidine ligands is the likely ancestral state. The lack of histidine ligands to Chl_D1_ and Chl_D2_ accounts for an up-shift in their E_*m*_ of +135 mV relative to the anoxygnic Type II reaction centres [40], which is consistent with the *oxidative jump* scenario. Even more remarkable is the observation that the E_*m*_ of P_D1_ in PSII is down-shifted by –135 mV indicating that the E_*m*_ of P has the potential to be substantial higher than +1200 mV. All in all, the structural shifts required to explain the divergence of Type I and Type II reaction centres may have resulted in a photosystem with an oxidising potential considerably above that required for the oxidation of water in Archean conditions. If this ancestral reaction centre was also capable of stabilising a primordial Mn/Ca cluster at the donor side, this would have led to a further increase in the E_m_ of P. In PSII the Mn_4_CaO_5_ cluster is calculated to contribute +200 mV to the E_m_ of P_D1_ [40].

After the helix shift, a tyrosine, hydrogen-bonded to a histidine and nearby Ca^2+^ binding site, could have become oxidised upon charge separation. This Tyr-His pair could have occupied a position similar to D468-Y513 in PshA and Y629-H601 in PsaB. The newly formed tyrosyl radical could have oxidised several aqueous Mn^2+^. Oxidised Mn may have been immediately stabilised by the Ca^2+^, a ligand from the antenna domain, and the carboxylic C-terminus, already present in the ancestral reaction centre as demonstrated here. Oxidation of Mn is followed by the ejection of a proton from any of the bound waters leading to the favourable formation of μ-oxo-bridges between adjacent Mn cations [41]. This process could have occurred in a manner very similar to the photoactivation of the Mn_4_CaO_5_ cluster in PSII, which can occur without the aid of chaperons or any specialised assembly factors [24, 60, 61].

## Final remarks

This speculative pathway represents a basic narrative for the structural changes that occurred at the divergence of Type I and Type II ancestral homodimeric reaction centres, but includes a key feature missing from previous models. This feature is an increase in the potential of the donor-side components and the acceptor-side components associated with a single event: a helix shift in the membrane. The proposed helix-shift induced change in redox potentials gave a selective advantage to the system by providing a strong oxidant that would have accessed new sources of electrons: most likely Mn followed by water. This structural shift may have resulted in a rapid cascade of changes and amino acid substitutions resulting in the optimisation of water oxidation, the establishment of linear electron transfer from water to ferredoxin, and the heterodimerisation of the core of PSII and PSI. Consistent with this perspective, we have recently shown that the gene duplication event leading to D1 and D2 is likely to have occurred soon after the Type I/Type II split, and before the L/M split of anoxygenic Type II reaction centres [18]. We have also shown evidence that these structural changes and duplication events were accompanied by rates of evolution of the reaction centre proteins at least 40 times greater than any rates observed in the past 2.5 billion years in any cyanobacterium, alga or plant. These fast rates indicate that water oxidation chemistry likely became optimised soon after the origin of the earliest reaction centres. The rate of evolution decreased exponentially and had stabilised at current levels before the most recent common ancestor of Cyanobacteria and well before the Archean/Proterozoic transition [18].

## Glossary

### Charge separation

Light-driven charge separation is the process of an electron in chlorophyll being excited to a higher energy level by the absorption of a photon. It is then transferred from the 1^st^ excited state orbital to a nearby electron acceptor thereby forming a radical cation and radical anion, the primary radical pair.

### Great Oxidation Event

Abbreviated as GOE. The time when the atmosphere became permanently oxygenated, about 2.4 billion years ago. Prior to the GOE and throughout all of Earth’s early history, it is thought that the atmosphere had a concentration of oxygen well below 10^-5^ of the present atmospheric level. However localised oxygen oases and “whiffs” of oxygen likely occurred for several hundred million years before the GOE.

### Mn_4_CaO_5_ cluster

The water-oxidising complex of Photosystem II. This is the catalytic site of Photosystem II where water oxidation occurs. It consists of 4 atoms of Mn, 1 Ca, and 5 bridging oxygens, arranged in a distorted-chair configuration.

### Most recent common ancestor

Also last common ancestor. The most recent common ancestor of any group of organisms is the most recent individual from which all the organisms in that group are directly descended. The most recent common ancestor of Cyanobacteria capable of oxygenic photosynthesis is the immediate ancestor of the genus *Gloeobacter*, the earliest branch in the tree, and of all other known species.

### Protocyanobacterium

A colloquial name historically used to refer to speculative ancestors of Cyanobacteria, which predate their most recent common ancestor by an undetermined amount of time.

### Photochemical reaction centres

Nature’s solar cells. These are protein complexes that convert the energy of light directly into chemical energy, through the movement of electrons via a series of redox cofactors, resulting in redox reactions, chemical bond formation, electric field formation and proton movements. The reaction centres can be of two types, referred to as Type I and Type II reaction centres. In Cyanobacteria and photosynthetic eukaryotes these are referred to as Photosystem I and Photosystem II respectively.

## Outstanding questions

When did water oxidation to oxygen originate?

How oxidising was the cationic radical form of the oxidising photochemical pigments in the ancestral reaction centre?

What evolutionary forces led to the divergence of Type I and Type II reaction centres? What is the function of the Ca^2+^-binding site in Heliobacteria?

Is there a Ca^2+^-binding site in other anoxygenic Type I reaction centres?

## Acknowledgements

The financial support of the Leverhulme Trust (RPG-2017-223) and the Biotechnology and Biological Sciences Research Council (BB/K002.627/1 and BB/L011206/1) is gracefully acknowledged

